# A calibration protocol for soil-crop models

**DOI:** 10.1101/2023.10.26.564162

**Authors:** Daniel Wallach, Samuel Buis, Diana-Maria Seserman, Taru Palosuo, Peter Thorburn, Henrike Mielenz, Eric Justes, Kurt-Christian Kersebaum, Benjamin Dumont, Marie Launay, Sabine Julia Seidel

## Abstract

Process-based soil-crop models are widely used in agronomic research. They are major tools for evaluating climate change impact on crop production. Multi-model simulation studies show a wide diversity of results among models, implying that simulation results are very uncertain. A major path to improving simulation results is to propose improved calibration practices that are widely applicable. This study proposes an innovative generic calibration protocol. The two major innovations concern the treatment of multiple output variables and the choice of parameters to estimate, both of which are based on standard statistical procedure adapted to the particularities of soil-crop models. The protocol performed well in a challenging artificial-data test. The protocol is formulated so as to be applicable to a wide range of models and data sets. If widely adopted, it could substantially reduce model error and inter-model variability, and thus increase confidence in soil-crop model simulations.

## 1 Introduction

Process-based models that describe crop growth and development, soil water and nitrogen dynamics and their interactions (henceforward “soil-crop” models) are an essential research tool for agronomy. They are the tool of choice for evaluating climate change impact on crop production and for testing adaptation and mitigation strategies (Asseng et al., 2019; Ramirez-Villegas et al., 2017; Webber et al., 2014). They are also used as aids in yield forecasting (van der Velde and Nisini, 2019), crop breeding programs (Ramirez-Villegas et al., 2020) and for informing crop management decisions (Keating et al., 2003).

A fairly recent practice, largely driven by the Agricultural Modeling Intercomparion and Improvement Project (AgMIP (Rosenzweig et al., 2013)) is to organize multi-model ensemble studies, where multiple modeling groups use the same inputs and simulate the same output variables. It has been observed that these studies systematically exhibit a large amount of variability between modeling groups (for example (Bruni et al., 2022; Webber et al., 2017), though this can be mitigated by using the multi-model mean or median (Martre et al., 2015; Wallach et al., 2018). In studies of the impact of global climate change on crop production using multiple soil-crop and multiple climate models, the contribution to total variability of variability among soil-crop models has been found to be even greater than the contribution due to variability in climate projections (Asseng et al., 2013; Li et al., 2015; Wang et al., 2020).

For given inputs, the variability in soil-crop model simulations arises from variability in the model equations (“model structure”) and in the values of the parameters. Comparing those two sources of uncertainty, it has generally been found that both are important. Uncertainty in model structure is often found to make the larger contribution to overall variability (Tao et al., 2018; Xiong et al., 2020; Zhang et al., 2017). However, a recent study pointed out that those previous studies assume that there is a fixed set of parameters to estimate, while in fact there is also considerable uncertainty in the choice of parameters to estimate. Taking into account the uncertainty in choice of parameters, it was found that parameter uncertainty in most cases contributed more, often much more, than model structure uncertainty to overall variability in simulations (Wallach et al., 2023a).

The necessity of improving model structure in order to reduce soil-crop model prediction error and inter-model variability has been recognized (Maiorano et al., 2017). Improving parameterization on the other hand, and in particular improving methodology of crop model calibration, has only recently been seen as a major pathway to improve predictions and reduce variability of soil-crop model simulations. In a series of multi-model studies, the AgMIP calibration group (https://agmip.org/crop-model-calibration-3/), considered the simplified situation where only phenology data were used for calibration and found that there was very substantial variability in calibration practices between modeling groups, even between modeling groups using the same model structure (Wallach et al., 2021a, 2021b, 2021c). The “role of users” on soil-crop model simulations has also been found elsewhere (Albanito et al., 2022; Confalonieri et al., 2016). The AgMIP group proposed a protocol for calibration based on phenology data, aimed at improving and homogenizing practices, and found that the new protocol substantially reduced variability and prediction error compared to the case where each modeling team implemented its usual calibration approach (Wallach et al., 2023b). This shows that it is possible to achieve the twin goals of reduced error and reduced variability of soil-crop model simulations through improved calibration practices.

The present study reports the next stage of the AgMIP calibration activity, which considers the general soil-crop model calibration problem, where one uses all available data and not just phenology data. The two objectives of this study are firstly to propose improved calibration practices for soil-crop models in order to reduce prediction error and secondly to formulate the recommendations in a protocol in such a way that they are applicable to essentially all soil-crop models and data sets, in order to reduce inter-modeling group variability.

Improved calibration practices are necessary in particular with respect to two problems. The first is how to take into account multiple variables. Data available for calibration may include multiple variables, such as days to several development stages, biomass, light interception and soil water at various dates, end of season measurements of yield, grain number and grain protein content and others. Furthermore, the variety of available data is increasing with new sensors, additional remote sensing possibilities and improved data transmission capabilities (Pasquel et al., 2022). It is important to take all observed variables into account, even for example if the focus of the study is only on yield, in order to simulate as realistically as possible the dynamics of the soil-plant system, since this should improve predictions for new environments (Angulo et al., 2013a; Pasley et al., 2023). More realistic simulations of all processes is also important if the model is used as a tool for understanding the contributions of different processes to an overall result

The major difficulty here is that as more variables are considered, the number of parameters to estimate increases, leading to numerical problems. The most common solution to this problem is to split the problem into parts by fitting the model to only one or only a few variables at a time, to avoid estimating a large number of parameters at the same time (Angulo et al., 2013b; Jha et al., 2021; Pasley et al., 2023). A variant of this approach is to also separately fit environments with and without water and nitrogen stresses (Guillaume et al., 2011; Kersebaum et al., 2011). It has been emphasized that it is important to choose the order so that the “most independent” variables are treated first (Pasley et al., 2023). The difficulty with this solution is that crop models describe an interacting system, and in general it is not possible to order the processes in such a way that earlier processes affect later processes but later processes have no effect on earlier processes; i.e. there are feedbacks in the system. As a result, when parameters of later processes are fit to data, this may degrade the fit to variables used in earlier calibration steps (Guillaume et al., 2011). There have not been any proposed procedures that allow one to fit all data simultaneously while nonetheless simplifying the numerical problem of finding the best parameter values.

The recommendations here propose doing the calibration calculations in two steps. In the first step, variables are treated individually. In the second step, the model is fit to all observed variables simultaneously, using weights based on the fit in the first phase. This is similar to the standard statistical approach of first doing ordinary least squares (OLS) regression, and then using the OLS results to obtain weights for doing weighted least squares (WLS) (Seber and Wild, 1989). This approach takes advantage of a particularity of soil-crop models, which is that while there are feedbacks, these are often quite limited. As a result, the first step is expected to give good staring points for searching for the best parameter values in the second step, thus greatly simplifying the numerical problem.

The second problem is how to choose the parameters to be estimated from the data. In general, crop models have many more parameters than can be estimated from available data, so it is necessary to decide which parameters to estimate. One approach identifies a priori (independently of the available data) the major parameters that should be estimated for a particular model structure (Laj R. Ahuja and Ma, 2011). Other studies have focused on ways of doing sensitivity analysis for crop models, in order to identify the most important parameters that should be estimated (Ceglar et al., 2011; Li and Ren, 2019). In some cases, selection involves an ad hoc combination of a priori choice, sensitivity analysis and test of various parameters to see how much they can improve the fit to the data. None of these approaches specifically addresses the problem of over-parameterization, or the risk of highly correlated and therefore highly uncertain parameter estimators. The AgMIP study on calibration using only phenology data proposed an original approach to choice of parameters to estimate, based on a standard model selection criterion (Wallach et al., 2023b). An analogous approach is used here, but extended to multiple measured variables. An alternative would be a Bayesian approach to parameter estimation, but this is rarely applied to soil-crop models (though see Dumont 2014 for a Bayesian approach to calibration of the STICS soil-crop model).

Previous calibration studies have very largely concerned calibration of a particular model(L.R. Ahuja and Ma, 2011). In order to reduce inter-model variability, which is the second objective of this study, it is necessary that the proposed calibration protocol be applicable to essentially any soil-crop model and to any set of measured data. To achieve this genericity, the protocol provides recommendations for procedures, and explains how to combine them with model expertise in order to apply those recommendations to each specific model. While much of the variability in simulations arises from the two problems discussed above, modeling teams may also differ as to other aspects of calibration (Wallach et al., 2021c). To further reduce variability therefore, the proposed protocol covers all the steps involved in soil-crop model calibration,

The proposed protocol was tested using the STICS soil-crop model (Beaudoin et al., 2023) with artificial data for winter wheat. Using artificial data makes it possible to evaluate prediction accuracy exactly, and to compare estimated and true parameter values. The “measured” data included days to three development stages, biomass at several dates, nitrogen content of final biomass, grain yield, grain protein and grain number. Altogether 23 parameters were considered. Both the number of different variables and the number of parameters considered are large compared to most studies. The protocol performed well including for out-of-sample predictions.

## 2. Methods

### 2.1. Structure of the artificial data set

In order to make the artificial data as realistic as possible, the data set is based on a real data set for winter wheat variety Apache grown in France, from variety trials carried out by Arvalis – Institut du vegetal Paris. The same environments (weather, soil characteristics, management) and the same measured variables as in that data set are used for the artificial data. The only difference is that the measured values are replaced by values simulated using the STICS model. The simulated variables are those shown in Table 1.

**Table 1.**
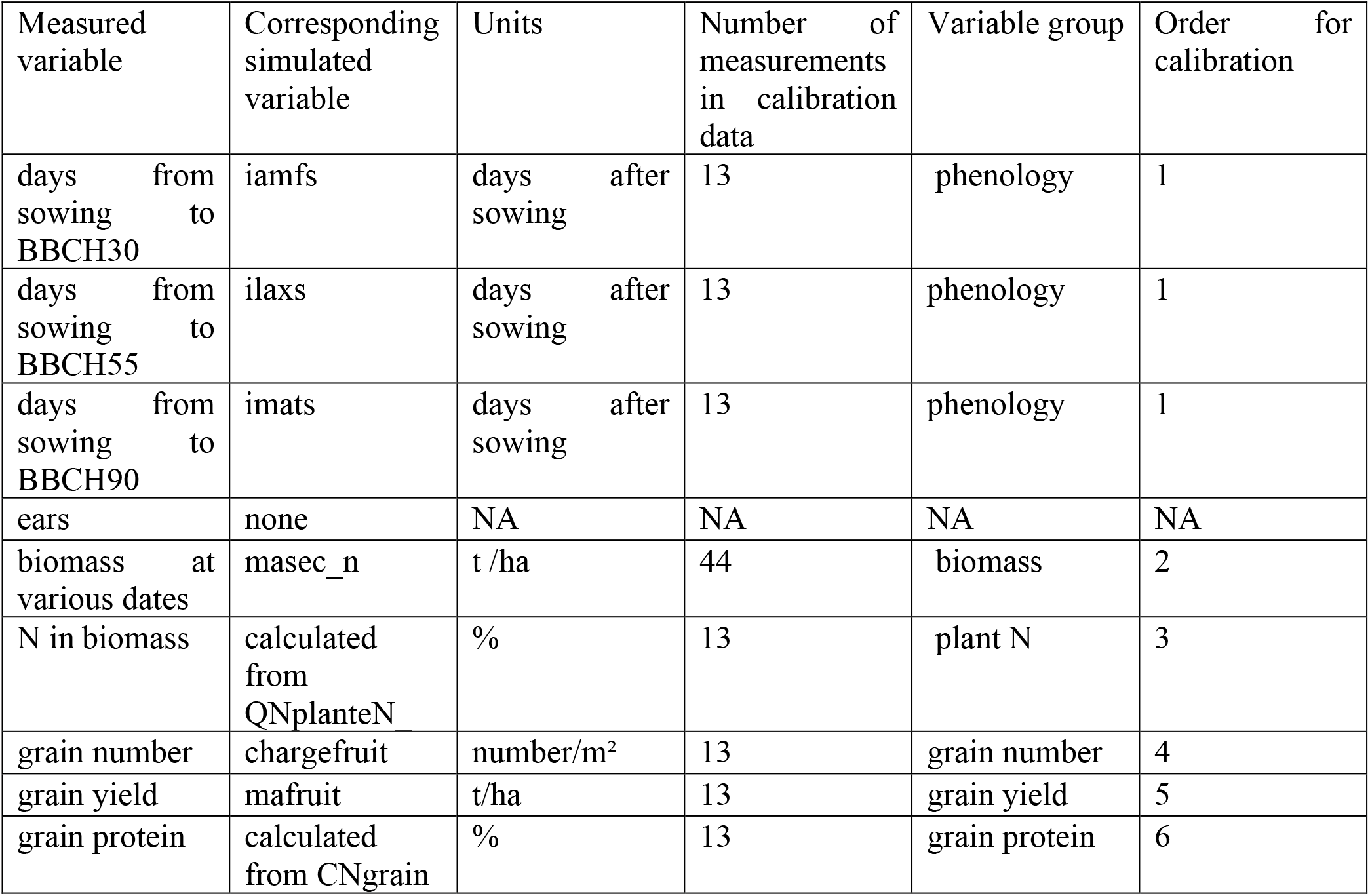
Measured and simulated variables, as an example of documentation for steps 2 and 3 of the protocol. There is one row for each measured variable, which also shows the corresponding simulated variable, if any, and the units of the simulated variable. The variables are grouped as explained in the protocol. The order of the groups is that in which the variable groups will be used for calibration. This example is for the STICS model applied to the artificial data for calibration used here. The development stages are stem elongation (BBCH30), heading (BBCH55) and maturity (BBCH90). Biomass refers to aboveground biomass.

| Measured variable | Corresponding simulated variable | Units | Number of measurements in calibration data | Variable group | Order for calibration |
| --- | --- | --- | --- | --- | --- |
| days from sowing to BBCH30 | iamfs | days after sowing | 13 | phenology | 1 |
| days from sowing to BBCH55 | ilaxs | days after sowing | 13 | phenology | 1 |
| days from sowing to BBCH90 | imats | days after sowing | 13 | phenology | 1 |
| ears | none | NA | NA | NA | NA |
| biomass at various dates | masec_n | t /ha | 44 | biomass | 2 |
| N in biomass | calculated from QNplanteN_ | % | 13 | plant N | 3 |
| grain number | chargefruit | number/m<sup>2</sup> | 13 | grain number | 4 |
| grain yield | mafruit | t/ha | 13 | grain yield | 5 |
| grain protein | calculated from CNgrain | % | 13 | grain protein | 6 |

The full data set has data from 22 environments, which were divided into two groups. The data from fourteen environments (six different sites, five different years) were used for calibration (the “calibration” data). The data from the eight other environments (five different sites, two different years) were used for testing (referred to as the “evaluation” or “out of sample” data). None of the sites or years present in the calibration data were also present in the evaluation data. Thus, the simulation errors for the evaluation data measure how well the calibrated model simulates for environments different than those used for calibration, but drawn from the same population (conventionally managed wheat fields in the major wheat growing regions of France, under current climate, sown with variety Apache), for the case where the data-generating mechanism is the same as for the calibration data. Further details about the environments can be found in Wallach et al. (2021a).

### 2.2 Generation of artificial data

The STICS soil-crop model, using parameter values previously estimated for the French winter wheat variety Soissons, was used to generate the artificial data for both the calibration and the evaluation environments. Those are henceforward referred to as the “true” parameter values. Then random noise was added to each generated value for the calibration environments. The amount of noise was chosen independently for each measurement by drawing from a Gaussian distribution with mean 0 and standard deviation equal to 2 days for the phenology data and equal to 10% of the generated value for the other variables, truncated at ± 3 standard deviations. Noise was not added to the evaluation data, since we are interested in prediction of the true values.

### 2.3 Default parameter values for the calibration

The parameter values used as starting values for the calibration (the “default” parameter values) of all the parameters shown in Tables 1 and 2 are different than their true values (Supplementary Table S3). The default values were chosen as the true values plus 60% of the distance to the assumed upper limit of the parameter or minus 60% of the distance to the assumed lower limit. The choice of whether to move the starting value toward the upper or lower limit was made at random independently for each parameter.

**Table 2.**
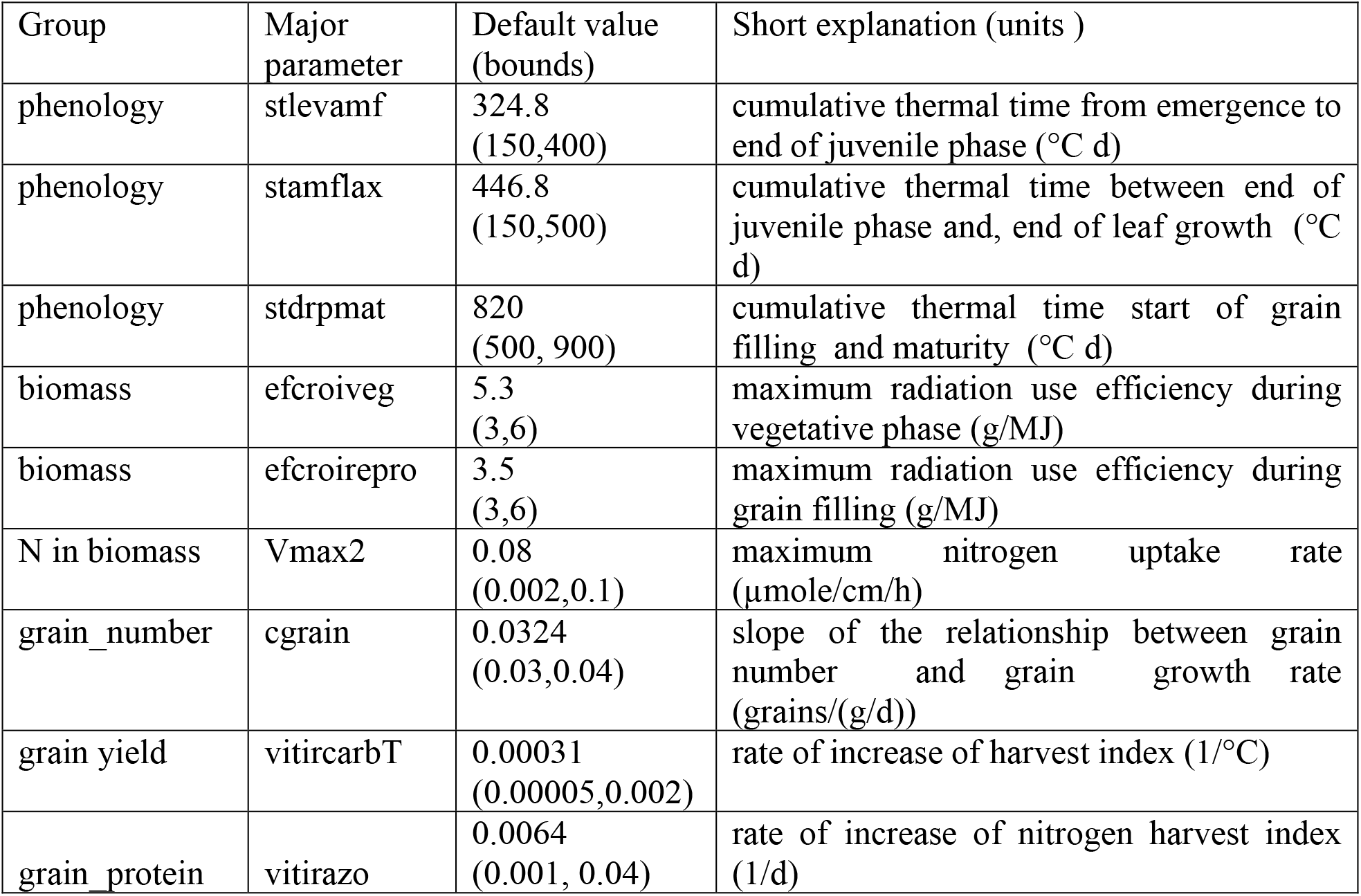
Major parameters. Example of documentation for step 4 of the protocol. There is one row for each major parameter of each variable group. The number of major parameters for each group is strictly limited, as explained in the protocol. The upper and/or lower bounds for each parameter can be specified. This example is for the STICS model applied to the artificial calibration data used here.

| Group | Major parameter | Default value (bounds) | Short explanation (units ) |
| --- | --- | --- | --- |
| phenology | stlevamf | 324.8<br>(150,400) | cumulative thermal time from emergence to end of juvenile phase (°C d) |
| phenology | stamflax | 446.8<br>(150,500) | cumulative thermal time between end of juvenile phase and, end of leaf growth (°C d) |
| phenology | stdrpmat | 820<br>(500, 900) | cumulative thermal time start of grain filling and maturity (°C d) |
| biomass | efcroiveg | 5.3<br>(3,6) | maximum radiation use efficiency during vegetative phase (g/MJ) |
| biomass | efcroirepro | 3.5<br>(3,6) | maximum radiation use efficiency during grain filling (g/MJ) |
| N in biomass | Vmax2 | 0.08<br>(0.002,0.1) | maximum nitrogen uptake rate (μmole/cm/h) |
| grain_number | cgrain | 0.0324<br>(0.03,0.04) | slope of the relationship between grain number and grain growth rate (grains/(g/d)) |
| grain yield | vitircarbT | 0.00031<br>(0.00005,0.002) | rate of increase of harvest index (1/°C) |
| grain_protein | vitirazo | 0.0064<br>(0.001, 0.04) | rate of increase of nitrogen harvest index (1/d) |

### 2.4 Calculations

For the test of the protocol, the R packages CroptimizR (Buis et al., 2023) and CroPlotR (Vezy et al., 2023) were used. All the calculations for steps 6 -8 were done automatically, using the tables prepared in steps 1-5 as input. A wrapper function for STICS available in the R package SticsOnR (Lecharpentier et al., 2023) was used to handle the communication between CroptimizR and the crop model (sending parameter values to the model and recovering simulated values). CroptimizR could be used with any crop model, but a different model wrapper would be required in each case.

The algorithm used in the example for searching the parameter space was the Nelder-Mead simplex algorithm (Nelder and Mead, 1965), as implemented in the R package nloptr R (Ypma and Johnson, 2022). This is a robust algorithm which is well-adapted to crop models since it does not use derivatives and does not require that the model be a continuous function of the parameters (Wang and Shoup, 2011). However, performance may become less efficient in high-dimension (Han and Neumann, 2006),The final result can depend on the starting values and the algorithm is not guaranteed to converge to a global minimum, so the implementation here used multiple starting points for the algorithm. For variable groups with more than one major parameter, the algorithm used 20 starting points within the upper and lower bounds of each major parameter, chosen by Latin Hypercube sampling. For variable groups with a single major parameter, five different starting points were used. For each candidate parameter, all previously chosen parameters had initial values equal to the optimal values previously found, and five different starting values were used for the new candidate. For step 7, one starting point was the best parameter values found when treating each variable group separately. In addition, 19 other starting points were generated at random using Latin Hypercube Sampling. In all cases, using the previous best values as starting point led to the lowest sum of squared errors.

### 2.5 Evaluation of the protocol

The following evaluation metrics were calculated

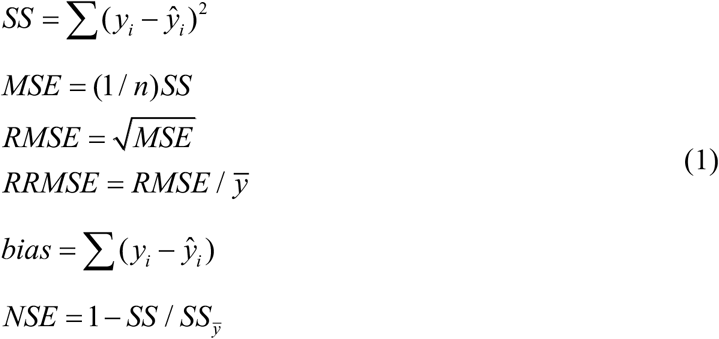

where MSE = mean squared error, RMSE = root mean squared error, RRMSE = relative root mean squared error, bias = model bias and NSE = Nash Sutcliffe efficiency. The sum in SS is over all measurements of the variable in question, *y*_*i*_ is the i^th^ measurement and 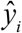 is the corresponding simulated value. In RRMSE, 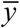 is the average of the measured values, and in NSE, 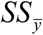 is the sum of squared errors for the model that uses 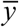 to predict for all environments. The above criteria are calculated with respect to the measured values, which includes measurement error, for the calibration data, and with respect to the true values, without measurement error, for the evaluation data.

## 2. Results

### 3.1 Description of the calibration protocol

The proposed protocol for soil-crop model calibration, described in detail below, is composed of eight steps (Figure 1). The first five steps (the model expertise part of the protocol) require detailed knowledge of the model and the data. No calculations are performed here. The result of these steps is a series of tables that contain all the model-specific information needed for the calculations. The last three steps (the calculation steps) describe the calculations to be done. The protocol includes instructions for each step, and the documentation to be produced in each step. The documentation is an integral part of the protocol, insuring transparency and reproducibility of the calibration procedure.

**Figure 1.**
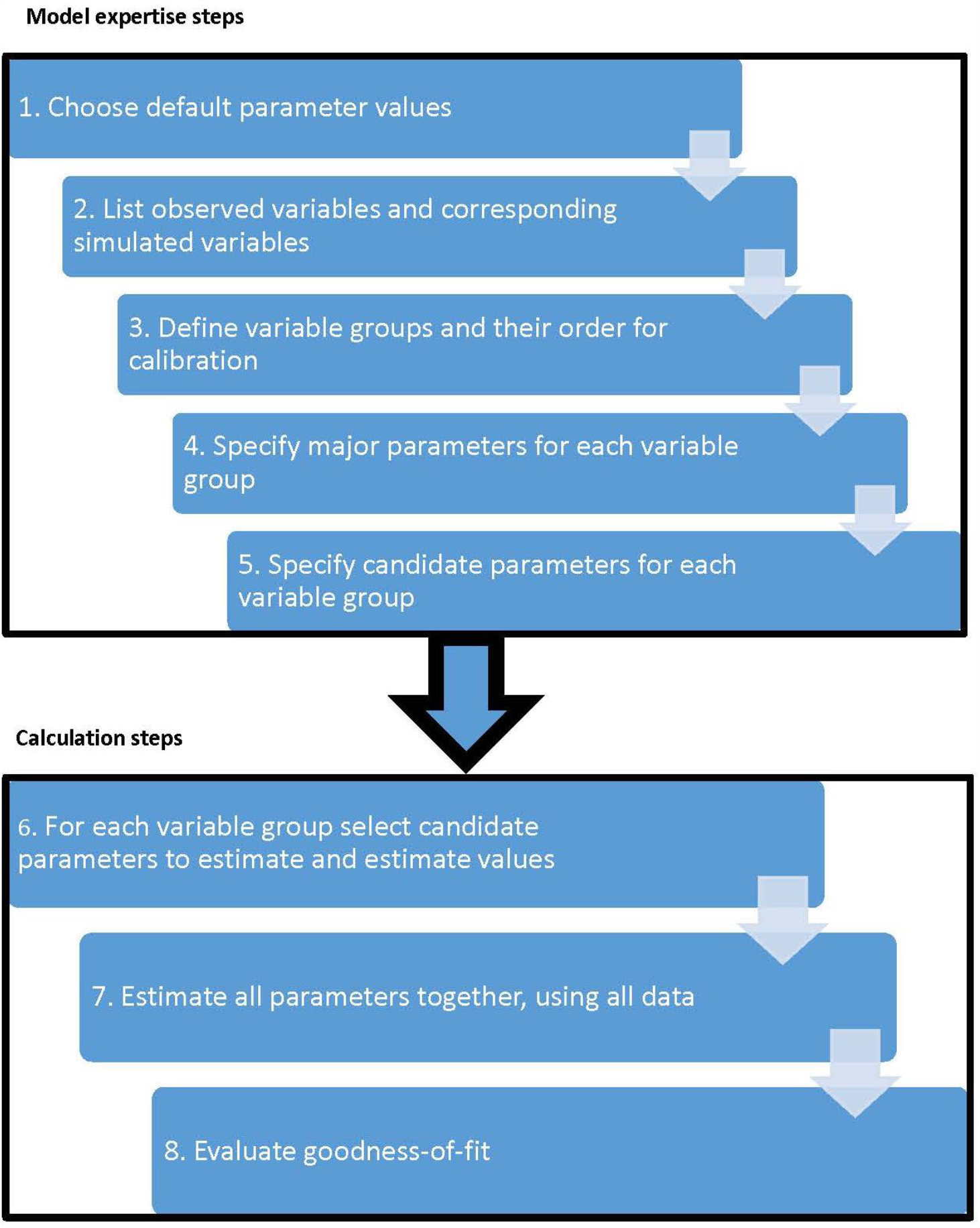
Schema of calibration protocol. The first five steps involve codifying model expertise. Given that information, the calculation steps 6 -8 require no farther model-specific inputs.

#### Step 1. Explain choice of default parameter values and describe the calibration environments

Only a small fraction of crop model parameters will usually be estimated from the data. The majority of the parameters will remain at their default values, so it is important to choose the default values with care. In particular, one should obtain as much information as possible about the cultivar characteristics (maturity class, photoperiod sensitivity, etc.) and choose default parameter values accordingly. The documentation required here (not shown) contains the cultivar characteristics and the rationale for the choice of default parameter values.

#### Step 2. List observed variables and corresponding simulated variables, if any

The purpose of this step is to identify the correspondence between observed and simulated variables. The documentation required for this step is a table with one row for each measured variable, showing also the corresponding simulated variable (Table 1 shows an example).

#### Step 3. Define groups of variables and order them

The grouping of observed variables is fixed by the protocol All days to development stages are grouped together in a phenology group. All measurements of a given variable at different times (e.g. biomass) will also be in the same group. Other variables (including all final values such as final yield, grain number, grain protein content etc.) will each constitute a separate group. The order of the groups is the order in which they will be used for fitting the model. The order of the groups is important. It should be such that the simulated values of the variables in a group have little or no effect on the simulated values of variables in previous groups. Phenology will usually be the first group, since the days to different development stages are usually not affected or only slightly affected by other variables. The required documentation here, which is combined with the documentation for step 2, shows the group and order for each observed variable (see Table 1).

#### Step 4. Identify the major parameter or parameters for each group of variables

The purpose of this step is to identify the major parameters that affect each variable group. There is a strict limit on the number of major parameters for each group, to avoid over-parameterization. If there is only one variable in the group, there can only be one major parameter. For variables with at least two measurements in some environments (e.g. biomass with in-season measurements) there can be at most two major parameters (for example, one that determines rate of increase during vegetative growth and a second that determines rate of increase during reproductive growth).. For phenology, there can be as many major parameters as observed development stages with simulated equivalents. However, each major parameter must affect the time to a different stage.

The major parameters for a group of variables should have an effect on the simulated values in all environments. If a parameter is nearly additive, i.e. has nearly the same effect in all environments, then the estimation of the parameter will make the model bias nearly zero for the associated variable, which is desirable. Thus, the first choice of major parameter for a variable is a parameter that is nearly additive. Thermal degree days to a development stage is usually a nearly additive parameter for days to that stage, since increasing the required number of degree days will, in general, increase the days to the stage by a similar amount for all environments. Parameters that describe the effect of stresses, which only affect the simulated values if the stresses are present, will not be major parameters. The required documentation here is a table which shows the major parameters for each variable group (see example in Table 2).

#### Step 5. Identify candidate parameters for each group of variables

The candidate parameters are those parameters that are likely to explain a substantial part of the variability between environments and/or management strategies that remains after the major parameters are estimated. Each of these parameters will be tested (in the next step), and will only be included in the final list of parameters to estimate if estimation leads to a sufficient improvement in fit to the data.

The candidate parameters should be ordered from supposedly most to supposedly least important. The number of candidate parameters is not limited, but it is recommended to keep the number fairly small. The required documentation here is a table with the candidate parameters for each variable group (see example in Table 3).

**Table 3.**
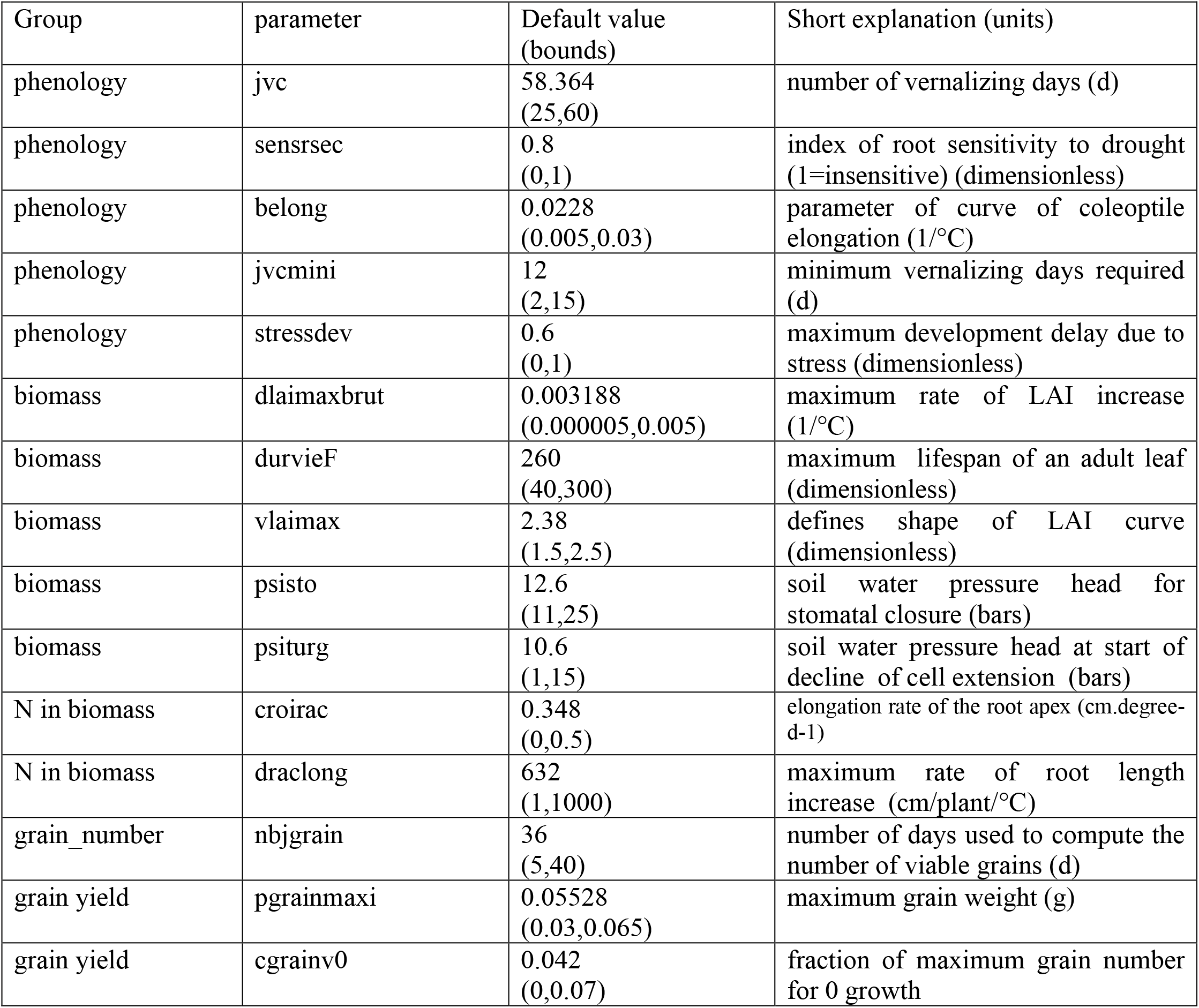
Candidate parameters. Example of documentation for step 5 of the protocol. There is one row for each candidate parameter for each variable group. The documentation includes the default value (i.e. best guess) of the parameter. Also, upper and/or lower limits for the parameter can be specified. This example is for the STICS model applied to the artificial calibration data used here.

#### Step 6. Selection of parameters to estimate for each variable group and first estimation of their values

In this step, each group of variables is treated separately, in the order chosen in step 3. A list of parameters to estimate for each group is initialized with the major parameters. The major parameters for the group are estimated using ordinary least squares (OLS), and the corrected Akaike Information Criterion (AICc Brewer et al., 2016; Chakrabarti and Ghosh, 2011) is calculated as

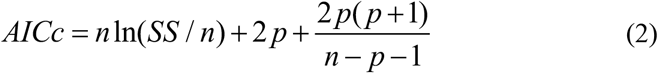

where SS is the sum of squared errors for all variables in the group, n is the number of data points and p the number of estimated parameters. This assumes that all model errors for the group are independent and identically normally distributed.

Once the major parameters have been estimated, each candidate parameter in turn is added tentatively to the list of parameters to be estimated. If estimating all the parameters on the list reduces AICc below the previous smallest value, the candidate is kept on the list of parameters to be estimated. Otherwise, the candidate is removed from the list of parameters to be estimated, and returns to its default value (see flow diagram in Figure 2). AICc is a standard model selection criterion, designed to choose the best predicting model even if none of the proposed models is the true model (Aho et al., 2014).

**Figure 2.**
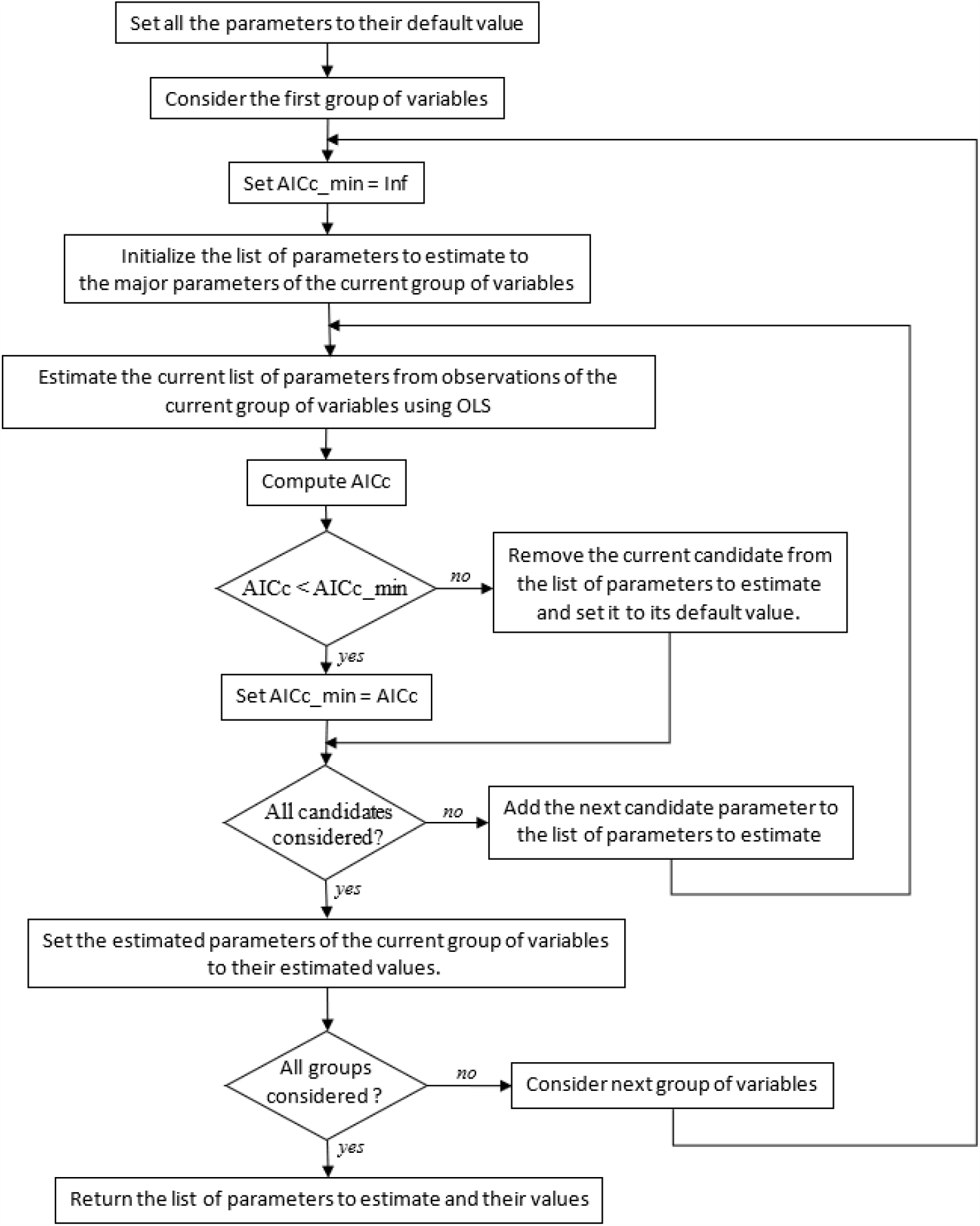
Flow diagram for the selection of parameters to estimate, and first estimation of their values, for one variable group. This is done in step 6 of the protocol

Biomass should be replaced by the natural logarithm of biomass before the calculation. The reason is that biomass values may go over a wide range of values during the growth period, with an associated increase in the standard deviation of model error. The log transformation will make the standard deviations approximately constant for all dates. Similarly, a log transformation should be used for any other variables expected to vary over a wide range over time.

In the example here, the Nelder-Mead simplex algorithm was used to find optimal parameter values, both in step 6 and step 7 (Nelder and Mead, 1965). However, other optimization algorithms could also be used. When testing candidate parameters, one of the starting points for optimization should be the previous best parameter values, since those will often be close to the new best values.

A first required documentation table here shows the result of adding each new candidate parameter, for each variable group (see example in Table 4). The optimum parameter values and the fit to the measured data after this step are combined with the documentation of step 7 (Table 5, Table 6).

**Table 4.** Selection of candidate parameters to estimate. Example of documentation for step 6 of the protocol. Each row shows the list of parameters to be estimated at that stage of the calculations. Candidate parameters that lead to a reduction in AICc compared to the previous smallest value are kept in the list of parameters to estimate. Otherwise, the candidate is removed from the list. This example is for the STICS model applied to the artificial calibration data used here, and for the variable group “biomass”. There will be an analogous table for each group of variables.

| Group | Parameters to be estimated | AICc | Candidate to be fit to data? |
| --- | --- | --- | --- |
| biomass | efcroiveg,efcroirepro | -127.89 | yes (automatically) |
| biomass | efcroiveg,efcroirepro<br>dlaimaxbrut | -166.13 | yes |
| biomass | efcroiveg,efcroirepro<br>dlaimaxbrut, durvieF | -164.97 | no |
| biomass | efcroiveg,efcroirepro<br>dlaimaxbrut, vlaimax | -178.70 | yes |
| biomass | efcroiveg,efcroirepro<br>dlaimaxbrut, vlaimax,psisto | -176.39 | no |
| biomass | efcroiveg,efcroirepro<br>dlaimaxbrut, vlaimax,psiturg | -177.02 | no |

**Table 5.** Parameter values. There is a row for each parameter that is estimated. This table is part of the documentation for steps 6 and 7 of the protocol. This example is for the STICS model applied to the artificial calibration data used here. The column of true values has been added here to facilitate evaluation of the protocol. In practical situations, the true values are unknown.

| Group | Estimated parameter | True value | Default value | Value after step 6 | Value after step 7 |
| --- | --- | --- | --- | --- | --- |
| phenology | stlevamf | 212 | 324.8 | 202.14 | 204.54 |
| phenology | stamflax | 367 | 446.8 | 356.63 | 358.95 |
| phenology | stdrpmat | 700 | 820 | 686.23 | 686.31 |
| phenology | belong | 0.012 | 0.0228 | 0.0061 | 0.0066 |
| phenology | stressdev | 0 | 0.6 | 0.087 | 0.093 |
| biomass | efcroiveg | 4.25 | 5.3 | 4.43 | 4.55 |
| biomass | efcroirepro | 4.25 | 3.5 | 3.91 | 3.43 |
| biomass | dlaimaxbrut | 0.00047 | 0.00318 | 0.00031 | 0.00032 |
| biomass | vlaimax | 2.2 | 2.38 | 2.08 | 2.09 |
| N in biomass | Vmax2 | 0.05 | 0.08 | 0.014 | 0.022 |
| grain_number | cgrain | 0.036 | 0.0324 | 0.037 | 0.035 |
| grain yield | vitircarbT | 0.0007 | 0.00031 | 0.00067 | 0.00068 |
| grain_protein | vitirazo | 0.0145 | 0.0064 | 0.015 | 0.014 |

**Table 6.** Relative root mean squared error (RRMSE) for the calibration data. The table shows RRMSE for the default parameter values and after parameter estimation in steps 6 and 7, for each variable. This table is part of the documentation for steps 6 and 7 of the protocol. This example is for the STICS model applied to the artificial calibration data used here.

|  | BBCH30 | BBCH55 | BBCH90 | ln(biomass) | N in biomass | grain number | grain yield | grain protein |
| --- | --- | --- | --- | --- | --- | --- | --- | --- |
| Default parameter values | 0.156 | 0.182 | 0.1588 | 0.051 | 0.22 | 0.31 | 0.25 | 0.286 |
| After step 6 | 0.013 | 0.013 | 0.0066 | 0.021 | 0.13 | 0.12 | 0.17 | 0.071 |
| After step 7 | 0.012 | 0.013 | 0.0065 | 0.018 | 0.12 | 0.11 | 0.16 | 0.076 |

#### Step 7. Re-estimation of all selected parameters using all variables simultaneously

In this step, all the selected parameters from step 6 are estimated together, using all the data, using weighted least squares (WLS). Each group of variables is weighted by the inverse of the standard deviation of model error (s), estimated from the results of step 6 as

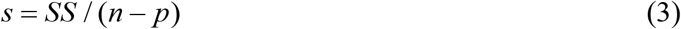

where SS is the sum of squared errors for all variables in the group, n is the number of data points and p the number of estimated parameters. The required documentation table here shows the estimated parameter values after steps 6 and 7 (see example in Table 5).

#### Step 8. Evaluation of goodness-of-fit

In this step metrics of goodness of fit are calculated for the simulations using the default parameter values, using the parameter values after step 6 and using the parameter values after step 7. The required documentation table here shows the metrics for goodness-of-fit at each stage. An example is shown in Table 6 and Supplementary Table S1.. Additional metrics could also be calculated. Graphs of simulated versus observed values for each variable should also be produced (see example in Supplementary Figures S1-S3).

### 3.2 Evaluation of the protocol

The STICS soil-crop model was used both to generate the artificial data and as the model calibrated using the protocol. The use of artificial data makes it possible to add measurement error as desired, rather than working with uncontrolled measurement error. It also makes it possible to compare estimated parameter values with the true values.

The six different variable groups in the artificial data are shown in Table 2. Data from fourteen environments were used for calibration, and data from 8 different environments were used to evaluate out-of-sample prediction error. For testing the protocol, 23 parameters of the STICS model were set to default values different than those used to generate the artificial data.

For all variable groups, the calibration substantially improved the fit to the calibration data (Table 6, Supplementary Table S1 and Figs. S1-S3). The improvement was most pronounced for the phenology group, where relative root mean squared error (RRMSE) decreased from 16%-18%, depending on the simulated development stage, using the default parameters to less than 1% for all stages using the parameters estimated in step 7. RRMSE for yield decreased from 25% with default parameter values to 16% after calibration., which was the largest final RRMSE value.

Calibration also substantially improved the fit to the out-of-sample data (Table 7, Supplementary Table S2). For the phenology group, RRMSE decreased from 14%-16% using the default parameters to less than 1% using the parameters estimated in step 7. RRMSE for yield decreased from 17% with default parameter values to 9% after calibration. The similar RRMSE values for the calibration and out-of-sample data show that there is not a problem of over-parameterization. For both calibration and out-of-sample data, RRMSE was smaller in almost every case after step 7 than after step 6, but the differences were in all cases small, at most a difference of 1.5% in RRMSE value. It seems that in this example feedbacks, which are ignored in step 6 where each variable group is fit separately, but which are taken into account in step 7; are relatively slight.

**Table 7.**
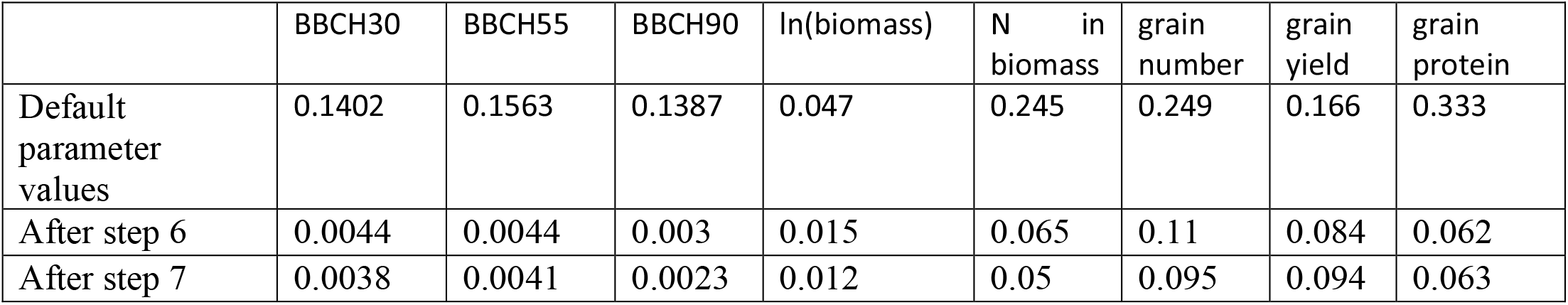
Relative root mean squared error (RRMSE) for simulation of out-of-sample data. The table shows RRMSE for the default parameter values and after parameter estimation in steps 6 and 7, for each variable. This example is for the STICS model applied to the artificial evaluation data used here.

All 23 parameters that had default values different than the values used to generate the artificial data were chosen as either major parameters or candidate parameters. However, only 13 of those parameters were finally estimated. The remainder were candidate parameters that did not reduce AICc, and so remained at their default values. Estimated values of the major parameters for the phenology group were much closer to the true values than were the default parameter values (Table 5). For other parameters the final estimated values could be closer or farther from the” true” values than were the default values. This suggests that there is relatively little compensation of errors for the major phenology parameters, because the major parameters have a preponderant effect on phenology, while the results for other variables depend on multiple parameters and so compensation of errors plays a larger role.

## 4. Discussion

The two objectives of the protocol are first to define calibration practices that are effective in reducing prediction error and secondly to reduce inter-model variability by formulating the protocol in such a way that it can be implemented by essentially all soil-crop models and for all data sets. The results here are very encouraging with respect to both objectives. The protocol was effective in substantially reducing errors both for the calibration environments and for out-of-sample environments compared to initial error, for all variables, despite the relatively large diversity of measured variables and the relatively large number of parameters that were considered. The protocol was in no way specifically adapted to the STICS model or to the artificial data set used here, so its successful application here suggests that it can be used as a standardized procedure for a large range of soil-crop models and data sets.

The use of artificial data for testing the protocol has both advantages and drawbacks. The major drawback is that the data are generated using the same model that is calibrated, which is never true in practice and which may artificially improve the performance of the protocol. This effect is mitigated by the fact that very many model parameters had starting values for calibration different than those used to generate the data, so that there are large differences between the data and the initial simulations. The advantage of artificial data is that one can compare the estimated parameters with the true parameter values, and the simulated responses with the true responses, including for out-of-sample environments.

It would be of interest to test alternative calibration procedures, using this protocol as a baseline. One alternative is to directly use all the data simultaneously for calibration, rather than first using one variable group at a time, despite the large number of parameters to estimate that that implies. One possibility here would be to use the PEST calibration package (Doherty et al., 2010), which uses regularization techniques to make the parameter estimation feasible even for highly correlated parameter estimators and for ill-conditioned models. There is still however the problem of choosing a limited number of parameters to consider. This might be done using sensitivity analysis (Necpálová et al., 2015).

Another alternative would be to do a Bayesian analysis, where one estimates the distribution of the parameters rather than the best value. In principle, the choice of which parameters to estimate is less crucial here than with a frequentist approach, since neither uninfluential nor highly correlated parameters create particular difficulties. However, even here it is not possible to consider all parameters, so some selection of parameters would still be necessary. Bayesian methods have so far been applied principally to calibration problems with relatively few variable types and parameters (Dumont et al., 2014; Iizumi et al., 2009; López-Cruz et al., 2017).

An important feature of the protocol here is the separation into two parts. The first part, based on model expertise, is where all the specificities of a particular model structure and data set are taken into account. The calculations in the second part can be completely automated, once the documentation from the first part is available. This could greatly simplify the calibration activity for soil-crop models. In the example here, all the calculations (steps 6-8) were done automatically, with the tables from steps 2-5 as inputs. The next step in the AgMIP calibration project is a multi-model application of the protocol to real data. This is currently underway.

## 5. Conclusions

Multi-modeling group simulation studies involving soil-crop models systematically result in a wide diversity of simulated results. This clearly limits the confidence one can have in the results and therefore their usefulness. There is no consensus as to best calibration practices, and it seems that the diversity in calibration approach is a major cause of variability among modeling groups. The protocol proposed here is a promising solution to the problem of calibration of soil-crop models. It has innovative solutions, based on statistical principles, to two major problems of crop-soil model calibration, namely the choice of parameters to estimate and the way to handle multiple outputs. Furthermore, it is applicable to a wide range of models and data sets. If widely adopted, this protocol could reduce errors and also reduce variability in simulated values between modeling groups compared to usual practice, and thereby improve the usefulness of soil-crop model simulations.

## Acknowledgements

This study was carried out in the framework of the Agricultural Model Intercomparison and Improvement Project (AgMIP). KCK was supported by AdAgriF - Advanced methods of greenhouse gases emission reduction and sequestration in agriculture and forest landscape for climate change mitigation (CZ.02.01.01/00/22_008/0004635). Partial funding was provided by the BonaRes project Soil3 (BOMA 03037514, 031B0515C) of the Federal Ministry of Education and Research (BMBF), Germany, and by Deutsche Forschungsgemeinschaft (DFG, German Research Foundation) under Germany’s Excellence Strategy-EXC 2070 – 390732324.

## Author contributions

DW was responsible for conceptualization, project administration and writing of the original draft. SB contributed to software and validation (adapted the CroptimizR software to automatically apply the protocol, and ran the software for the STICS model). DMS contributed to validation (tested the software using the model HERMES). SB, SS, TP, PT and HM contributed to project administration. EJ, CK, BD and ML contributed to validation (provided model expertise for the STICS or HERMES models). All authors contributed to conceptualization and review and editing. All authors read and approved the final manuscript.

## Data availability

All the results of this study can be reproduced based on input data and artificial measured data, using the STICS model. The model is publicly available. The input and artificial measured data are available from the authors on reasonable request.

