## supplementary for "A calibration protocol for soil-crop models"

Table S1

Multiple goodness-of-fit metrics for the calibration data. The metrics are mean squared error (RMSE), relative root mean squared error (RRMSE), Nash-Sutcliffe Efficiency (NSE) and model bias (bias). The table shows results for the default parameter values and after parameter estimation in steps 6 and 7, for each variable. This table is part of the documentation for steps 6 and 7 of the protocol. This example is for the STICS model applied to the artificial calibration data used here.

|  |  | BBCH30 | BBCH55 | BBCH90 | ln(biomass) | N in biomass | grain_number | grain yield | grain_protein |
| --- | --- | --- | --- | --- | --- | --- | --- | --- | --- |
| RMSE | default | 24.1 | 37.5 | 41.1 | 0.32 | 0.24 | 6474 | 186 | 2.3 |
|  | step6 | 1.9 | 2.7 | 1.7 | 0.13 | 0.14 | 2444 | 128 | 0.57 |
|  | step7 | 1.8 | 2.7 | 1.7 | 0.11 | 0.13 | 2305 | 124 | 0.61 |
| RRMSE | default | 0.156 | 0.182 | 0.1588 | 0.051 | 0.22 | 0.31 | 0.25 | 0.286 |
|  | step6 | 0.013 | 0.013 | 0.0066 | 0.021 | 0.13 | 0.12 | 0.17 | 0.071 |
|  | step7 | 0.012 | 0.013 | 0.0065 | 0.018 | 0.12 | 0.11 | 0.16 | 0.076 |
| NSE | default | -4.21 | -16.5 | -31.78 | 0.92 | -1.9688 | -1.88 | 0.5 | -2.36 |
|  | step6 | 0.97 | 0.91 | 0.94 | 0.99 | 0.0014 | 0.59 | 0.76 | 0.79 |
|  | step7 | 0.97 | 0.91 | 0.94 | 0.99 | 0.1513 | 0.63 | 0.78 | 0.76 |
| bias | default | -23.5 | -37.36 | -41 | -0.209 | 0.175 | -656 | 107.4 | 1.9537 |
|  | step6 | 0.214 | -0.71 | -0.5 | 0.036 | 0.07 | 178 | -1.6 | -0.015 |
|  | step7 | 0.071 | -1.07 | -0.71 | 0.016 | 0.037 | -381 | 15.6 | -0.0129 |

Table S2

Multiple prediction error metrics for the evaluation data. The metrics are mean squared error (RMSE), relative root mean squared error (RRMSE), Nash-Sutcliffe Efficiency (NSE) and model bias (bias). The table shows results for the default parameter values and after parameter estimation in steps 6 and 7, for each variable. This example is for the STICS model applied to the artificial evaluation data used here.

|  |  | BBCH30 | BBCH55 | BBCH90 | ln(biomass) | N in biomass | grain_number | grain yield | grain_protein |
| --- | --- | --- | --- | --- | --- | --- | --- | --- | --- |
| RMSE | default | 22.64 | 33.17 | 36.53 | 0.294 | 0.274 | 5433 | 135 | 2.63 |
|  | step6 | 0.71 | 0.94 | 0.79 | 0.095 | 0.072 | 2388 | 68 | 0.49 |
|  | step7 | 0.61 | 0.87 | 0.61 | 0.075 | 0.056 | 2072 | 76 | 0.5 |
| RRMSE | default | 0.1402 | 0.1563 | 0.1387 | 0.047 | 0.245 | 0.249 | 0.166 | 0.333 |
|  | step6 | 0.0044 | 0.0044 | 0.003 | 0.015 | 0.065 | 0.11 | 0.084 | 0.062 |
|  | step7 | 0.0038 | 0.0041 | 0.0023 | 0.012 | 0.05 | 0.095 | 0.094 | 0.063 |
| NSE | default | -3 | -14.35 | -20.8 | 0.93 | -11.78 | -4.92 | -1.03 | -10.82 |
|  | step6 | 1 | 0.99 | 0.99 | 0.99 | 0.11 | -0.14 | 0.48 | 0.59 |
|  | step7 | 1 | 0.99 | 0.99 | 1 | 0.47 | 0.14 | 0.36 | 0.58 |
| bias | default | -21.88 | -33 | -36.38 | -0.197 | 0.267 | -5054 | 108 | 2.58 |
|  | step6 | 0.25 | 0.62 | 0.62 | 0.041 | 0.056 | 891 | 31 | -0.21 |
|  | step7 | 0.12 | 0 | 0.38 | 0.017 | 0.026 | 87 | 49 | -0.21 |

Table S3

Parameters with default values different than true values. List of parameters of the STICS model with default values for calibration different than the true parameter values (i.e. Than the values used to generate the artificial data). For each parameter, the variable group that it mainly affects is indicated.

| Parameter | Variable group mainly affected | Short explanation | True value | Default value |
| --- | --- | --- | --- | --- |
| stlevamf | phenology | cumulative thermal time emergence) and end of juvenile phase (°C d) | 212 | 324.8 |
| stamflax | phenology | cumulative thermal time between end of juvenile phase and, end of leaf growth (°C d) | 367 | 446.8 |
| stdrpmat | phenology | cumulative thermal time between start of grain filling and maturity (°C d) | 700 | 820 |
| jvc | phenology | number of vernalizing days (d) | 55.91 | 58.364 |
| sensrsec | phenology | index of root sensitivity to drought (1=insensitive) (dimensionless) | 0.5 | 0.8 |
| belong | phenology | parameter of curve of coleoptile elongation (1/°C) | 0.012 | 0.0228 |
| jvcmini | phenology | minimum vernalizing days required (d) | 7 | 11 |
| stressdev | phenology | maximum development delay due to stress (dimensionless) | 0 | 0­.6 |
| efcroiveg | biomass | maximum radiation use efficiency during vegetative phase (g/MJ) | 4.25 | 5.3 |
| efcroirepro | biomass | maximum radiation use efficiency during grain filling (g/MJ) | 4.25 | 3.5 |
| dlaimaxbrut | biomass | maximum rate of LAI increase (1/°C) | 0.00047 | 0.003188 |
| durvieF | biomass | maximum lifespan of an adult) (dimensionless) | 200 | 260 |
| vlaimax | biomass | defines shape of LAI curve (dimensionless) | 2.2 | 2.38 |
| psisto | biomass | soil water pressure head for stomatal closure (bars) | 15 | 12.6 |
| psiturg | biomass | soil water pressure head at start of decline of cell extension (bars) | 4 | 10.6 |
| Vmax2 | N in biomass | maximum nitrogen uptake rate (µmole/cm/h) | 0.05 | 0.08 |
| croirac | N in biomass | elongation rate of the root apex (cm.degree-d-1) | 0.12 | 0.348 |
| draclong | N in biomass | maximum rate of root length increase (cm/plant/°C) | 80 | 632 |
| cgrain | grain number | slope of the relationship between grain number andgrain growth rate (grains/(g/d)) | 0.036 | 00324 |
| nbjgrain | grain number | number of days used to compute the number of viable grains (d) | 30 | 36 |
| vitircarbT | grain yield | rate of increase of harvest index (1/°C) | 0.0007 | 00.00031 |
| pgrainmaxi | grain yield | maximum grain weight | 0.0407 | 0.05528 |
| cgrainv0 | grain yield | fraction of maximal grain number for 0 growth | 0 | 0 |
| vitirazo | grain protein | rate of increase of nitrogen harvest index (1/d) | 0.0145 | 0.0064 |

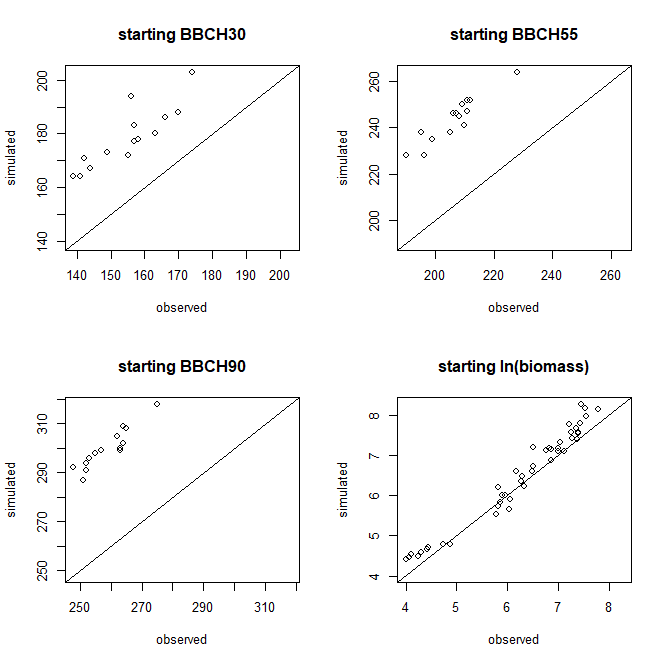

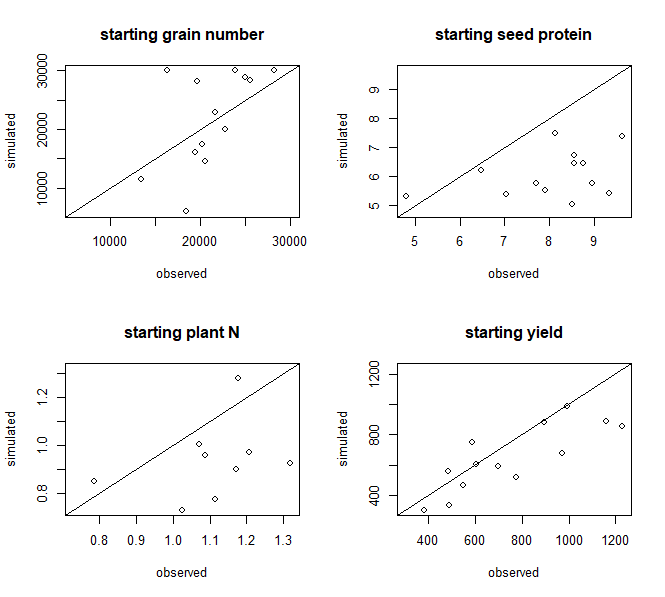

Figure S1

Simulated versus observed values, for the calibration data, using the STICS model with default parameter values.

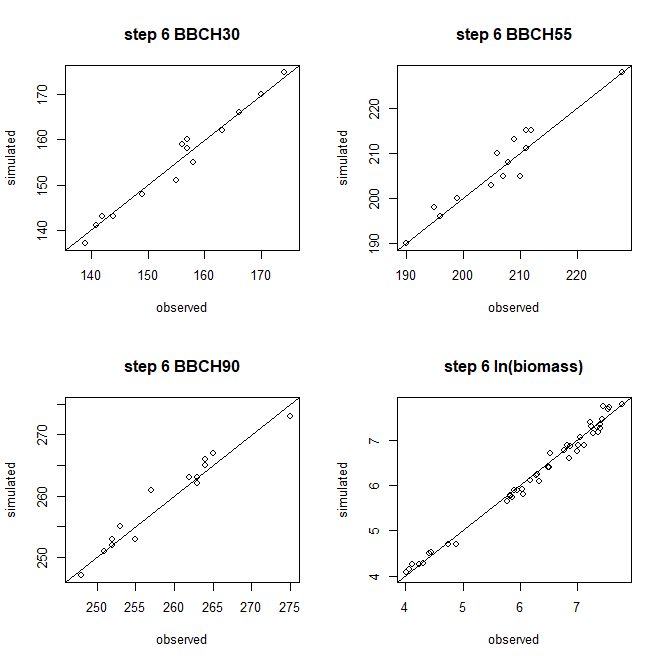

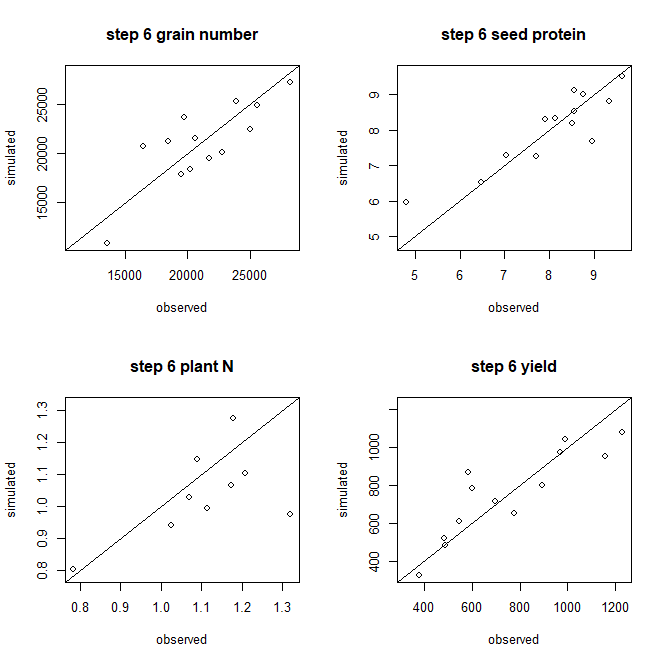

Figure S2

Simulated versus observed values, for the calibration data, using the STICS model with parameter values estimated in step 6 of the calibration protocol.

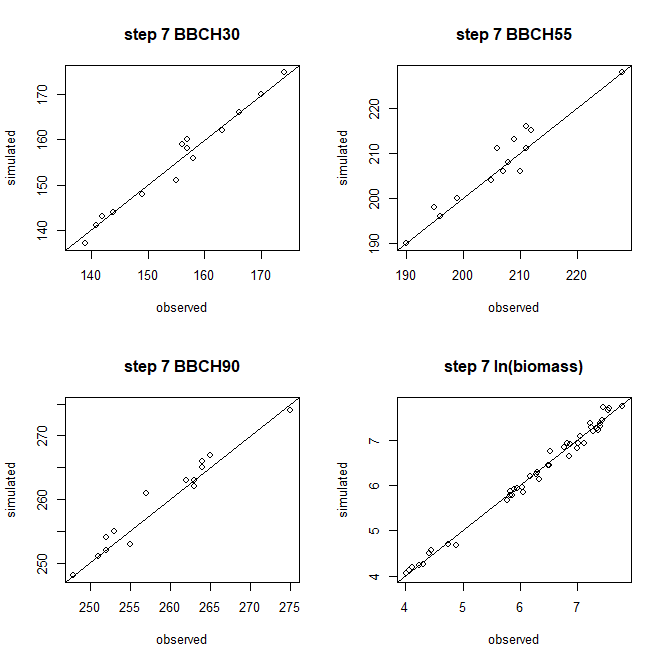

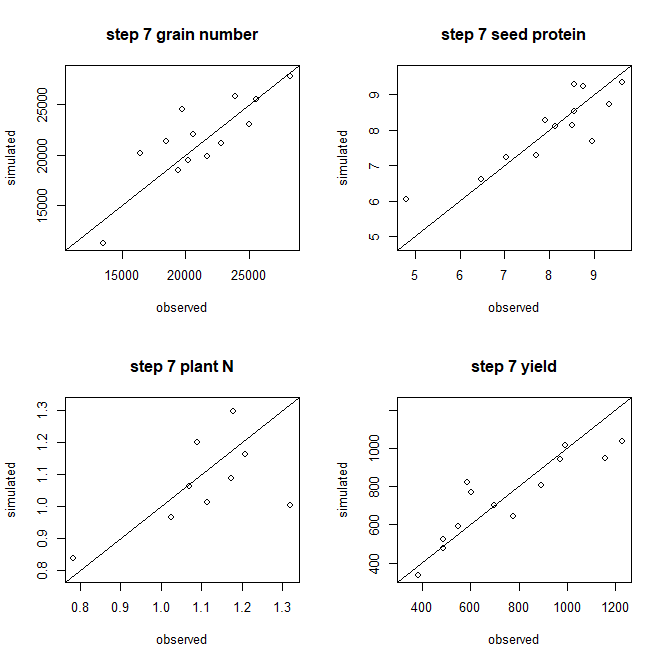

Figure S3

Simulated versus observed values, for the calibration data, using the STICS model with parameter values estimated in step 7 of the calibration protocol.
